# Inhibition of vaccinia virus L1 *N*-myristoylation by the host *N*-myristoyltransferase inhibitor IMP-1088 generates non-infectious virions defective in cell entry

**DOI:** 10.1101/2022.06.13.495866

**Authors:** Lalita Priyamvada, Wouter W. Kallemeijn, Monica Faronato, Kimberly Wilkins, Cynthia S. Goldsmith, Catherine A. Cotter, Suany Ojeda, Roberto Solari, Bernard Moss, Edward W. Tate, Panayampalli Subbian Satheshkumar

**Affiliations:** Poxvirus and Rabies Branch, Centers for Disease Control and Prevention, Atlanta, GA 30329. USA; Department of Chemistry, Molecular Sciences Research Hub, Imperial College London, London W12 0BZ, UK; The Francis Crick Institute, 1 Midland Road, London NW1 1AT, United Kingdom; Infectious Diseases Pathology Branch, Centers for Disease Control and Prevention, Atlanta, GA 30329. USA; Laboratory of Viral Diseases, National Institute of Allergy and Infectious Diseases, National Institutes of Health, Bethesda, MD 20892, USA; National Heart and Lung Institute, Imperial College of Science, Technology & Medicine, London W2 1PG, UK

**Author notes:** Correspondence (E.W.T.), (P.S.S.). Equal contribution (Lalita Priyamvada, Wouter W. Kallemeijn). Clinipace, Morrisville, NC 27560, USA.

**Keywords:** Vaccinia virus, *N*-myristoylation, *N*-myristoyltransferases (NMTs), IMP-1088, NMT inhibitor, virus entry, quantitative chemical proteomics, poxvirus, antivirals

## Abstract

We have recently shown that the replication of rhinovirus, poliovirus and foot-and-mouth disease virus requires the co-translational *N-*myristoylation of viral proteins by human host cell *N*-myristoyltransferases (NMTs), and is inhibited by treatment with IMP-1088, an ultrapotent small molecule NMT inhibitor. Here, we reveal the role of *N*-myristoylation during vaccinia virus (VACV) infection in human host cells and demonstrate the anti-poxviral effects of IMP-1088. *N-*myristoylated proteins from VACV and the host were metabolically labelled with myristic acid alkyne during infection using quantitative chemical proteomics. We identified VACV proteins A16, G9 and L1 to be *N-*myristoylated. Treatment with NMT inhibitor IMP-1088 potently abrogated VACV infection, while VACV gene expression, DNA replication, morphogenesis and EV formation remained unaffected. Importantly, we observed that loss of *N*-myristoylation resulted in greatly reduced infectivity of assembled mature virus particles, characterized by significantly reduced host cell entry and a decline in membrane fusion activity of progeny virus. While the *N*-myristoylation of VACV entry proteins L1, A16 and G9 was inhibited by IMP-1088, mutational and genetic studies demonstrated that the *N*-myristoylation of L1 was the most critical for VACV entry. Given the significant genetic identity between VACV, monkeypox virus and variola virus L1 homologs, our data provides a basis for further investigating the role of *N*-myristoylation in poxviral infections as well as the potential of selective NMT inhibitors like IMP-1088 as broad-spectrum poxvirus inhibitors.

## INTRODUCTION

Vaccinia virus (VACV), the protype member of the family *Poxviridae* and genus orthopoxvirus (OPXV) is closely related to variola virus (VARV), the causative agent of smallpox. VACV-induced immunity through smallpox vaccination cross*-*protected against VARV and led to its eradication [1]. The remaining VARV stocks and materials are consolidated and stored in two high containment laboratories for studies aimed to improve diagnostics, determine vaccine efficacies, and develop new antivirals [2]. Due to the discontinuation of smallpox vaccination programs and waning immunity, a vast majority of the global population is immunologically naïve to smallpox and will require intervention by vaccination or antivirals for pre-and post-exposure therapeutics in the event of an exposure. To this end, we are interested in exploring the importance of post-translational modifications (PTMs) of viral proteins for the viral life cycle, and whether interference in PTMs is a potential antiviral strategy against OPXV.

*N*-myristoylation, a lipidic co-and post-translational modification of many eukaryotic proteins, involves the attachment of myristic acid, a 14-carbon saturated fatty acid, to the glycine (G) residue at the protein N-terminus [3-5]. The addition of the myristic acid moiety is catalyzed by *N*-myristoyltransferase enzymes (NMTs) and occurs after the removal of the initiator methionine (M) residue from the N-terminal “MG” motif by methionine aminopeptidases [4]. In humans, NMT exists in two isoforms, NMT1 and NMT2, which are highly conserved across all mammals [6]. The *N*-myristoylation of proteins by NMTs can have different effects on the substrate such as membrane anchoring, which in turn facilitates cellular processes involving protein localization, protein-protein interactions and signaling [7-9]. Recently, unbiased chemical biology approaches using a myristic acid analog (YnMyr, tetradec-13-ynoic acid) for metabolic incorporation, bio-orthogonal modification and enrichment by pulldown followed by mass spectrometry were successful in identifying the global cellular content of *N-*myristoylated proteins in human cells [10-12]. The viability of several pathogens, including various parasites and viruses, has been shown to be dependent on protein *N*-myristoylation. While parasites like *Plasmodium* and *Trypanosoma* encode their own NMT, host NMTs are usurped by viruses [13]. *N*-myristoylation of viral proteins is observed or predicted in families ranging from RNA to nucleo-cytoplasmic large DNA viruses, including *Picornovirdae, Arenaviridae, Reoviridae, Retroviridae, Hepadnaviridae, Polyomaviridae, Ascoviridae, Herpesviridae, Poxviridae, Asfiviridae* and *Iridoviridae* [13].

Previously, we identified and characterized small molecule inhibitors against *Plasmodium, Leishmania* and *Trypanosoma* NMTs [14-17], which were subsequently developed into inhibitors with high specificity and potency against human NMTs [18]. We further showed that one of these small molecules, IMP-1088, a sub-nanomolar EC_50_ dual inhibitor of human NMT1 and NMT2, blocked rhinovirus replication at low nanomolar concentrations [19]. The loss of *N*-myristoylation induced by IMP-1088 hampered viral replication and virus*-*induced cell death by blocking viral capsid assembly while causing minimal cytotoxicity in multiple rhinovirus strains, poliovirus and foot-and-mouth disease virus [19]. To determine whether the antiviral activity of IMP-1088 extends beyond picornaviruses to other viruses that require *N*-myristoylation, we evaluated its *in vitro* efficacy against VACV.

The VACV genome encodes over 200 open reading frames, including 11 proteins with a putative *N*-myristoylation motif (MG, at the N-terminus). Using radio-labelled myristic acid, four proteins (A16, E7, G9 and L1) were previously determined to undergo *N*-myristoylation [20]. Three of these proteins (A16, G9 and L1) are membrane proteins that associate and form the entry/fusion complex (EFC) required for virus entry [21]. While *N*-myristoylation was not required for membrane localization of L1, mutation of the glycine to an alanine in the N-terminal “MG” motif rendered VACV non-infectious [22]. Additional VACV proteins with predicted *N*-myristoylation motifs are involved in functions ranging from morphogenesis (membrane formation), DNA repair and genome formation, to virus spread and host range factors (Supplementary Table 1). Given the wide range of functions associated with the putatively *N-*myristoylated VACV proteins, we explored whether the intracellular inhibition of host NMT by IMP-1088 affects VACV replication.

The objectives of this study were to: 1) identify host and viral *N-*myristoylated proteins during VACV infection by quantitative chemical proteomics, 2) assess the impact of IMP-1088 on host and viral protein *N*-myristoylation, and 3) evaluate the inhibitory effect of IMP-1088 on VACV infection. Our results demonstrate that IMP-1088 potently inhibits VACV infection and spread with minimal cytotoxicity *in vitro*. We find that the VACV proteins A16, G9 and L1 undergo *N*-myristoylation, with L1 most strongly and significantly responding to NMT inhibition by IMP-1088. Moreover, loss of *N*-myristoylation resulted in the generation of entry-defective VACV virions while not significantly affecting viral gene expression, DNA replication, morphogenesis and viral egress. Taken together, our data demonstrate that blocking the host cell driven *N*-myristoylation of VACV protein L1 is a potent anti-VACV antiviral strategy.

## RESULTS

### IMP-1088 inhibits VACV spread and virus yield

To determine the impact of NMT inhibitor IMP-1088 treatment on VACV infection, we quantified virus yield, spread, and cellular cytotoxicity at different IMP-1088 concentrations upon infection of HeLa cells with VACV WR-GFP (0.3 multiplicity of infection, MOI) as illustrated in Figure 1A. A reduction of >50% in both virus yield (Figure 1B) and viral spread (Figure 1C) was measured at approximately 100 nM, indicating that IMP-1088 is a potent inhibitor of VACV infection. No cellular toxicity was observed in the presence of up to 10 μM IMP-1088, a concentration nearly 100-fold higher than its EC_50_, yielding a selectivity index >100 (Figure 1D).

**Figure 1.**
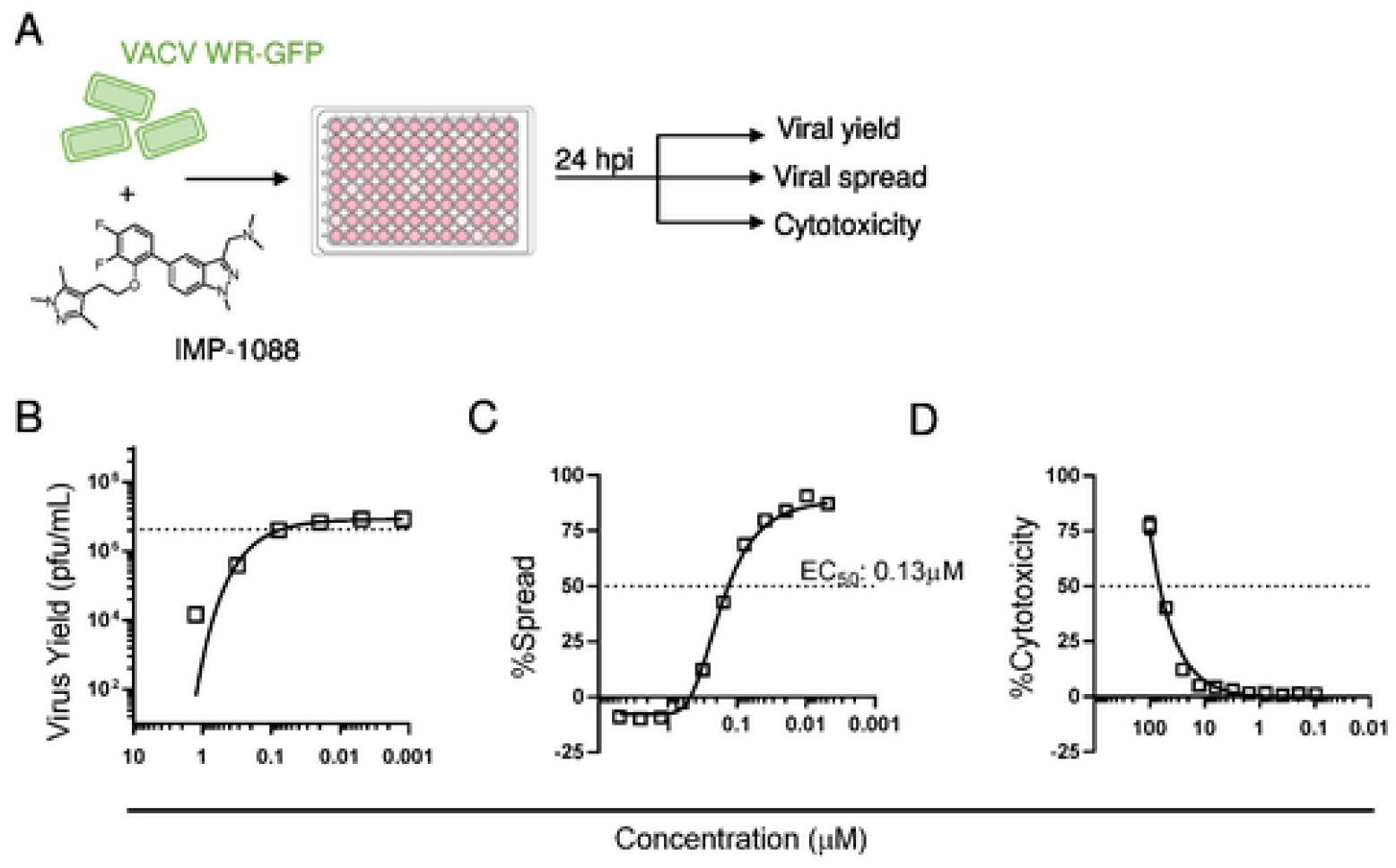
IMP-1088 inhibits VACV spread and virus yield. (A) Step-wise illustration of virus yield, spread and cytotoxicity assays. (B) Quantification of VACV yield in the presence of IMP-1088. HeLa cells were infected with VACV WR at increasing concentrations of IMP-1088. The cells were harvested 24 hpi, lysed by freeze-thaw and virus yield was determined by plaque assay. Dotted line indicates 50% virus yield. (C) Measuring viral spread in the presence of IMP-1088 based on GFP expression. Cells were infected with VACV WR-GFP in the presence of different concentrations of IMP-1088. The concentration of IMP-1088 required to reduce viral spread by 50% (EC_50_) was determined. (D) Cytotoxic effects of IMP-1088 determined by LDH assay. The concentration of IMP-1088 required for 50% of maximum OD (LDH signal in the absence of inhibitor) was measured and is reported as CC50. All experiments were performed twice with two replicates in each experiment. Values represent means +/-SEM.

### Quantitative chemical proteomics demonstrate inhibition of host and VACV L1 *N*-myristoylation by IMP-1088

Metabolic labelling with YnMyr, an alkyne-tagged myristic acid analog, allows the identification and quantification of *N-*myristoylated proteins encoded in the proteome of both the host and VACV, and provides quantitative insights induced by blocking NMT activity via IMP-1088.

In a multiparameter chemical proteomics experiment, we examined the effect of VACV infection on proteins enriched following YnMyr labelling, and in the presence or absence of NMT inhibitor IMP-1088. As shown in Figure 2A, after robust statistical testing of the chemical proteomics data, we identified 115 VACV proteins (out of 220), of which 3 are significantly enriched after YnMyr labelling: A16, G9 and L1. Analysis of the VACV proteome, retrieved from UniProt, revealed that VACV may express 11 proteins with a glycine at the second position (after the initiator methionine, see Supplementary Table 1), including the proteins A16, G9 and L1 identified by chemical proteomics. This evidence suggests A16, G9 and L1 are likely to be *N-*myristoylated.

**Figure 2.**
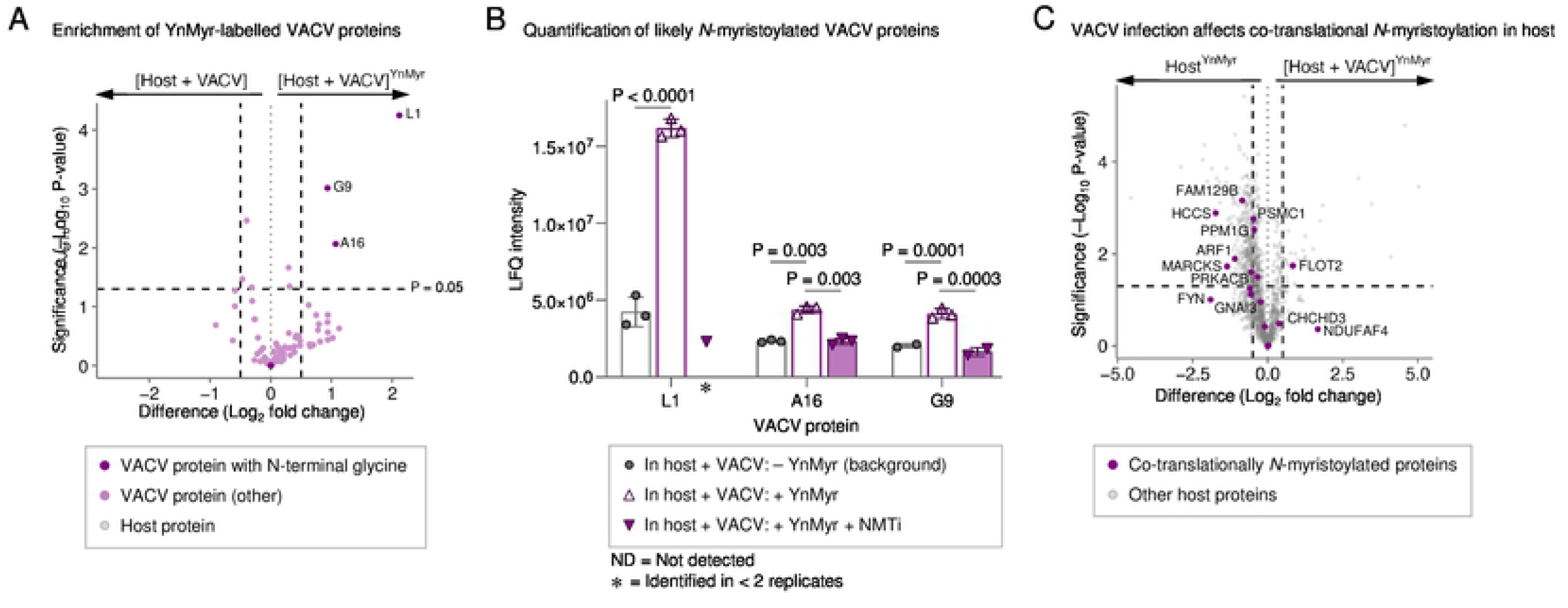
Chemical proteomic identification and quantification of host and viral proteins after VACV infection, enrichment, and mass spectrometric detection. (A) Identification of VACV proteins after YnMyr enrichment. Left vertical line depicts -0.5 Log_2_ fold change, right of +0.5; horizontal line depicts significance cut-off (P = 0.05); 3 *N*-myristoylated VACV proteins depicted in purple, 115 VACV proteins in pink, 2738 human proteins in gray. (B) Label free quantification intensity of *N-*myristoylated VACV proteins L1, A16, G9, as determined in background, after metabolic tagging with YnMyr and after NMT inhibition with IMP-1088. Average of 3 replicates, error bars depict standard deviation, significance tested by ANOVA. (C) Effect of VACV infection on 32 known co-translationally *N-*myristoylated protein levels of the host (purple). Left vertical line depicts -0.5 Log_2_ fold change, right of +0.5; horizontal line depicts significance cut-off (P = 0.05). Other proteins in gray.

Given the VACV proteome may contain additional putative NMT substrates, we analyzed the reduction of YnMyr labelling in the presence of NMT inhibitor IMP-1088 (Figure 2B and Supplementary Figure 1). In-depth analysis of label-free quantification (LFQ) intensities revealed that YnMyr-dependent enrichment of A16, G9 and L1 is depleted to background levels upon loss of *N*-myristoylation due to NMT inhibition (Figure 2B). Statistical significance could not be calculated for the change in L1 enrichment upon NMT inhibition, as the peptide levels reduced below the detection limit in two of three replicates, suggesting L1 is an NMT substrate with particularly high sensitivity to inhibition of NMT activity (Figure 2B). Moreover, YnMyr enrichment of A16 and G9 reduces most significantly upon NMT inhibition compared to other VACV proteins (Figure 2B and Supplementary Figure 1), suggesting this is specifically due to a loss of *N*-myristoylation. The observed reduction of *N*-myristoylation supports the hypothesis that VACV proteins require host NMTs, as there is no evidence of a VACV-encoded NMT. Target engagement of IMP-1088 towards the host NMTs, in the absence of VACV infection, was validated by the identification and significant depletion of YnMyr-dependent enrichment of 32 known co-translationally *N-*myristoylated substrates (Supplementary Figure 2). Next, we investigated the effect of VACV infection on the levels of proteins metabolically tagged with YnMyr, both in terms of known co-translationally and post-translationally *N-*myristoylated NMT substrates. As shown in Figure 2C, VACV infection reduces the levels of a majority of the 32 known co-translational *N-*myristoylated proteins that were identified by chemical proteomics. Similar reductions are seen with 27 known post-translational *N-*myristoylated proteins (Supplementary Figure 3). This observation corresponds with earlier findings that VACV infection reduces the synthesis of host proteins, in favor of proteins required for progressing the VACV life cycle, and thereby also likely affecting the flux of NMT substrates [23, 24].

### Host NMT inhibition does not affect VACV gene expression and morphogenesis

The impact of host NMT inhibition by IMP-1088 was analyzed at different steps in the VACV life cycle. The effect of IMP-1088 on gene expression or DNA replication was tested using VACV WR expressing luciferase under an early/late and late promoter, respectively (Figure 3A). We found no significant difference in early protein synthesis at 2 hours post infection (hpi) between cells infected in the presence or absence of IMP-1088 (Figure 3B). At 24 hpi, the presence of IMP-1088 also caused no observable change in late protein synthesis, suggesting that IMP-1088 did not impact VACV DNA replication (Figure 3C). Late protein expression requires VACV DNA replication as demonstrated by inhibition with the DNA replication inhibitor AraC (Figure 3C). Additionally, we observed multiple morphological forms of virus that are generated during infection, namely crescent (C), immature virus (IV), mature virus (MV), wrapped virus (WV) and enveloped virus (EV), in both untreated and IMP-1088 treated cells via electron microscopy (Figure 3D), demonstrating no qualitative effect on VACV morphogenesis and virion formation.

**Figure 3.**
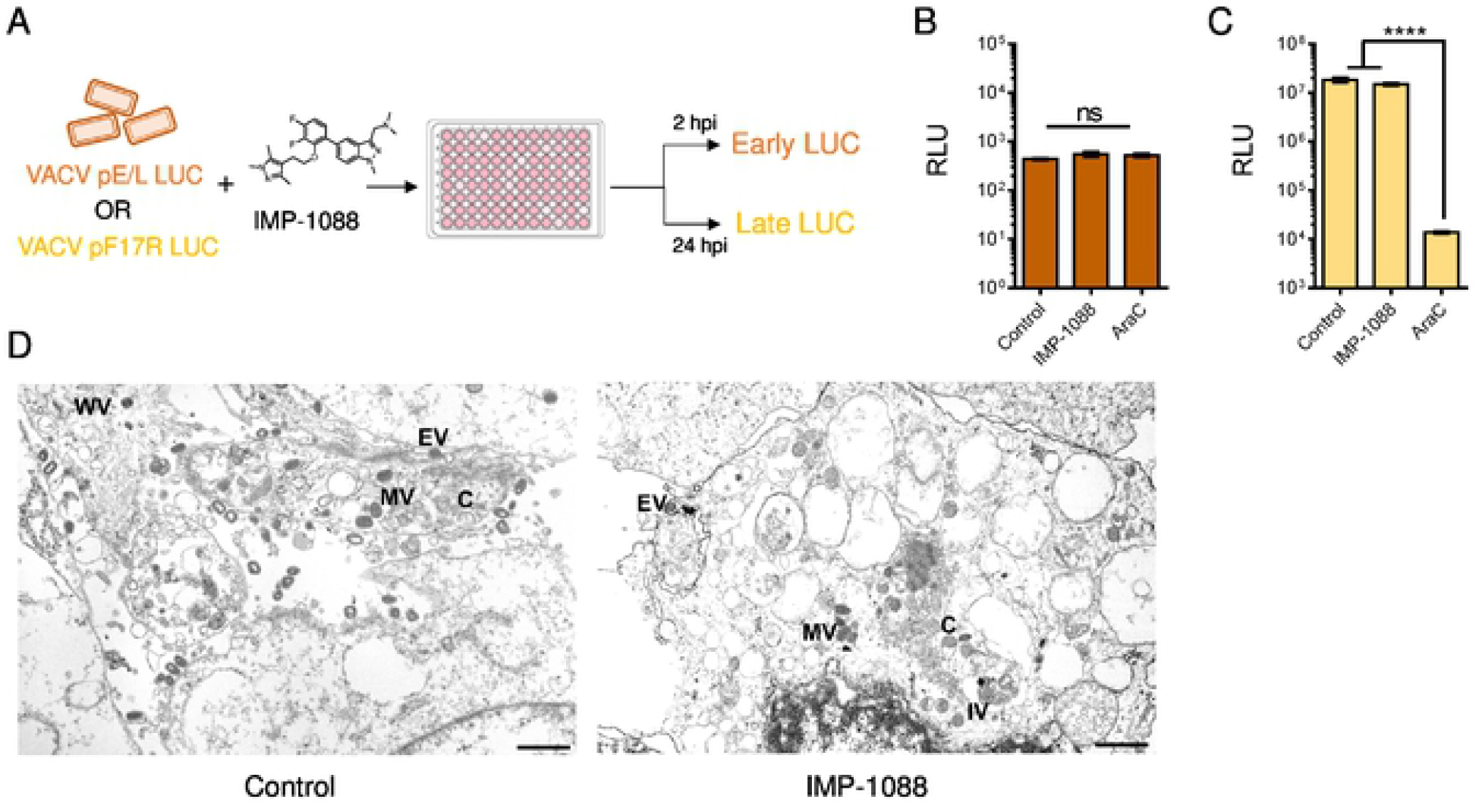
IMP-1088 does not affect VACV gene expression and morphogenesis. (A) Schematic representation of early and late protein detection in the presence of 2 μM IMP-1088. (B) BSC40 cells were infected with purified VACV WR-pE/L LUC virus in the absence and presence of 2 μM IMP-1088 and 40 μg/mL AraC. The level of secreted luciferase from early promoter was determined at 2 hpi. (C) BSC40 cells were infected with purified VACV WR-pF17R LUC in the absence and presence of 2 μM IMP-1088 and 40 μg/mL AraC. Luciferase levels were measured 24 hpi. RLU values of virus control, IMP-1088 and AraC treatments in (B) and (C) were compared using a one-way ANOVA followed by a Tukey’s multiple comparisons test. Ns indicates no significant difference and **** signifies a *p<0*.*0001*. (C) Transmission electron micrographs of VACV-infected cells in the absence and presence of IMP-1088. The various morphogenic forms of VACV are seen in both treatments; crescent (C), immature virus (IV), mature virus (MV), wrapped virus (WV) and extracellular virus (EV). Scale bar corresponds to 1 μm.

### Host NMT inhibition decreases infectivity of progeny VACV particles

Given the lack of an apparent effect of IMP-1088 on viral replication, membrane wrapping or assembly, we queried whether virions generated in the presence of IMP-1088 were non-infectious. The yield of MV purified from VACV infected cells treated with IMP-1088 (Figure 4A, lanes 2 and 4) was around 2-fold lower than that of control untreated cells as determined by protein staining (Figure 4A, lanes 3 and 5). When similar amounts of MV purified in the absence and presence of IMP-1088 were subjected to DNA isolation and tested for encapsidated genomic DNA by quantitative PCR (qPCR), similar Ct values were obtained at multiple dilutions (Figure 4B). Based on the qPCR results, equivalent numbers of viral particles from control and IMP-1088 treated cells were tested for infectious virus yield by plaque assay (measured in plaque forming units, pfu). Compared to untreated cells, MV purified from IMP-1088 treated cells exhibited a nearly 4-log reduction in infectivity (Figure 4C). This very high particle to pfu ratio was characteristic of virions with an entry-deficient phenotype.

**Figure 4.**
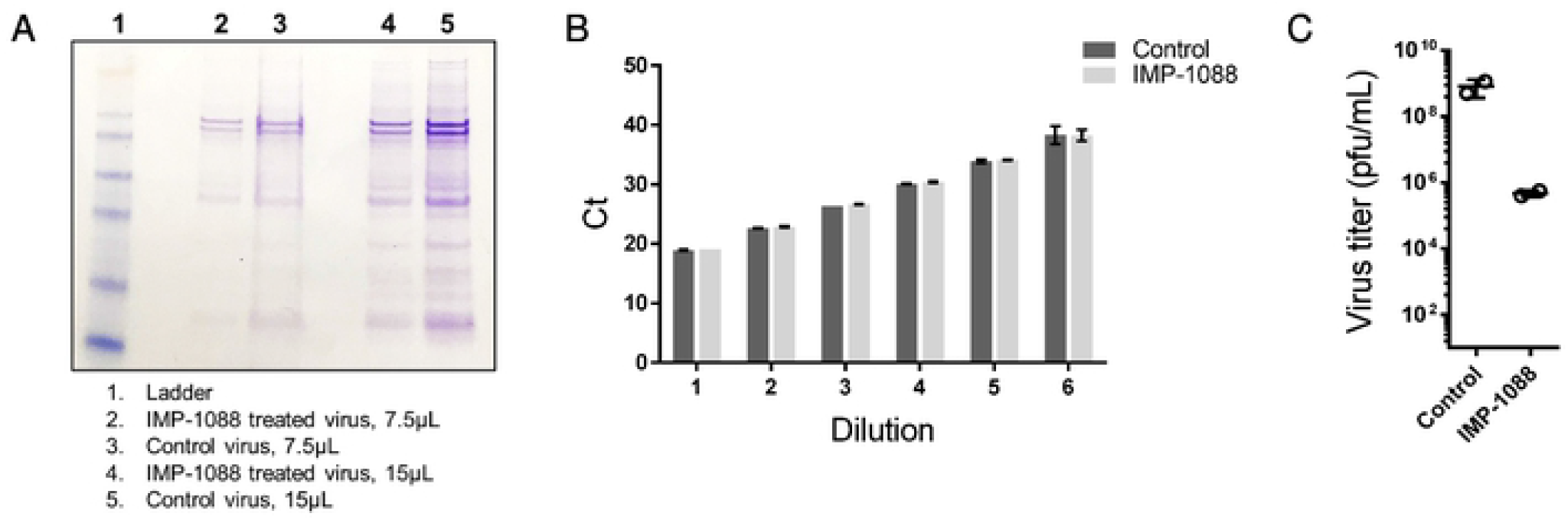
IMP-1088 decreases infectivity of progeny VACV particles. (A) Coomassie stained SDS-PAGE gel showing difference in protein levels between IMP-1088-treated and untreated (control) viruses. BSC40 cells were infected with VACV WR pE/L-LUC for 24 h in the absence and presence of 2 μM IMP-1088. Cells were harvested and lysed, and virus particles were purified by sucrose density gradient centrifugation. Either 7.5 or 15 μl of purified virus was run in a 4-12% SDS PAGE and stained with Coomassie blue. (B) Equivalent viral particles based on Coomassie staining were subjected to DNA isolation followed by real time PCR using VACV-specific primers. Ct values at different dilutions of purified DNA from the two treatments are shown. (C) Equivalent viral particles from untreated and IMP-1088 treated virus were tested by plaque assay to determine virus yields. Yields plotted for both treatments as pfu/mL.

### Host NMT inhibition reduces EV infectivity without affecting yield

During VACV infection, a large majority of virus particles generated are of the MV type, a small proportion (1-10%) of which undergo an additional double-membrane wrapping step to form EV particles. We queried whether treatment of virus with IMP-1088 affected EV yield. The VACV strain IHD-J was used instead of WR because of the higher number of EV particles released. EV particles released in the cell culture medium were collected to measure viral yield and to quantitate the number of viral particles as depicted in Figure 5A. Since entry of EV into cells is dependent on the EFC, we anticipated that EVs produced in the presence of IMP-1088 would have low or no infectivity. Indeed, infectious virus yield in the presence of IMP-1088 was lower than that of ST-246 treated cells (an inhibitor of EV formation), with values similar to AraC, which prevents genome replication (Figure 5B). In the absence of infectivity, we quantified the number of secreted EV particles by qPCR following DNA isolation. Only a slight reduction in viral genomic DNA content was observed with IMP-1088 treatment compared to untreated virus (Figure 5C). However, the reduction in Ct values was much more pronounced with AraC or with ST-246. Taken together, these data demonstrate that IMP-1088 reduces the infectivity of both MVs and EVs without significantly impacting virus particle formation and egress.

**Figure 5.**
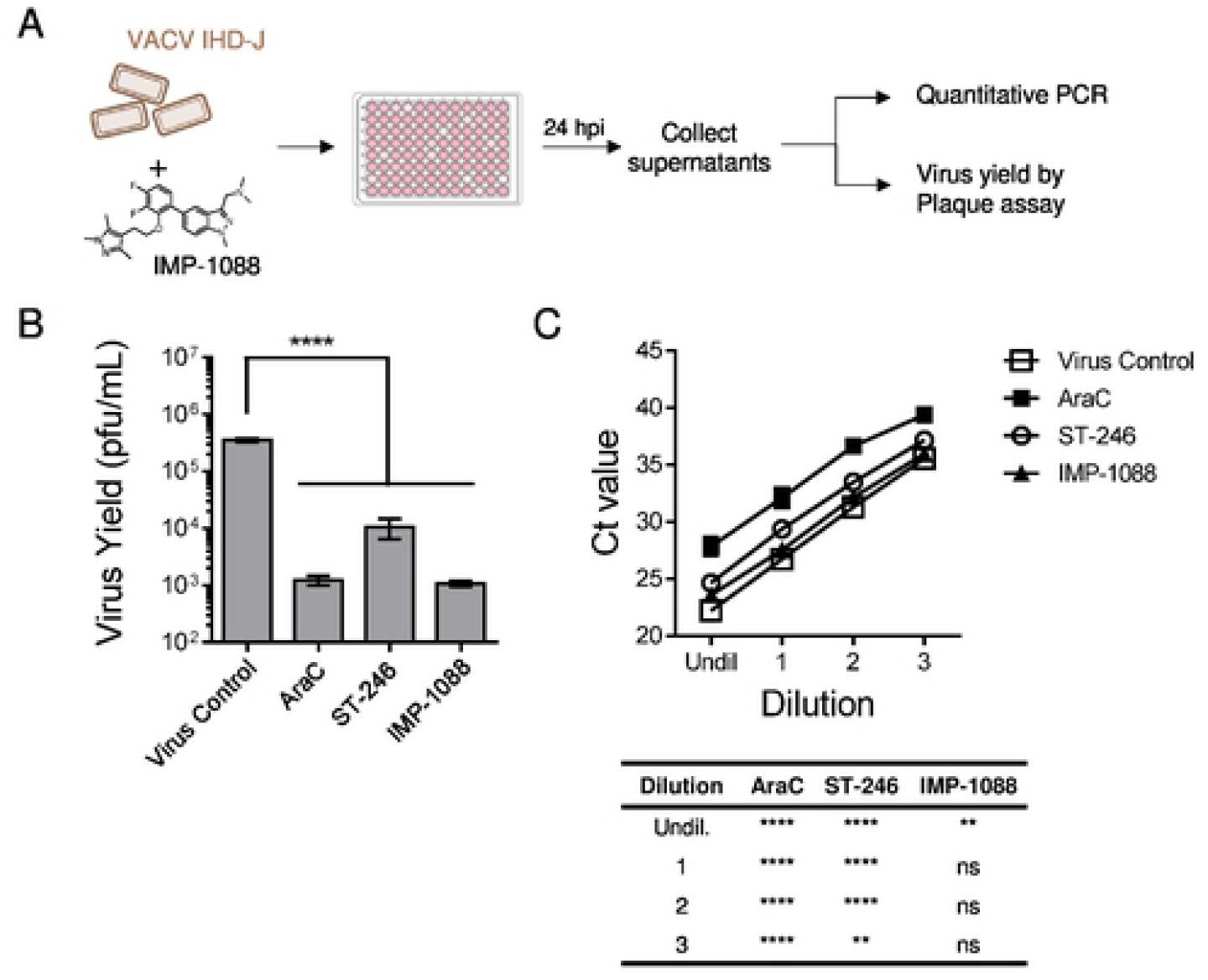
IMP-1088 reduces EV infectivity without affecting yield. (A) Schematic representation of assays used to assess VACV EV production after treatment with IMP-1088. (B) BSC40 cells were infected with VACV IHD-J in presence of inhibitors AraC, ST-246 and IMP-1088 for 24 h. The culture media was collected, spun at low speed to remove debris and cells, and tested by plaque assay to determine virus yield (pfu/mL). Three replicates were tested per viral dilution for each treatment, and the mean values +/-SD are shown. (C) Total viral DNA in the culture media for all four treatments was quantified by DNA isolation followed by qPCR. Four replicates per dilution were tested for every treatment, and the means +/-SD are shown. In both B and C, a one-way ANOVA was performed to determine statistical significance, followed by Tukey’s multiple comparisons test. Ns= not significant, ** = p≤0.005 and ****=p<0.0001.

### Decrease in VACV infectivity under host NMT inhibition is linked to a defect in viral entry

To measure the impact of IMP-1088 on virus entry, cells were infected with VACV WR-pE/L-LUC virus purified from cells with or without IMP-1088 treatment for 2 h and the level of early gene expression was determined by luciferase assay. The results demonstrated a statistically significant (p<0.0005) difference in luciferase levels between the virus purified from untreated versus IMP-1088 treated cells, with the latter largely remaining below the limit of detection (Figure 6A). The data are consistent with a delay in the release of viral cores into the cytoplasm of cells, which is required to trigger early gene expression. We also examined membrane fusion by infecting cells labeled with a lipophilic dye, DiO and determined dye transfer by flow cytometry. Although not statistically significant, there was an observable reduction in the frequency of DiO+ cells incubated with virus purified from IMP-1088 treated cells compared to the untreated control, indicating reduction in VACV hemifusion with the host cell membrane (Figure 6B). Laliberte et al (2011) found that the absence of L1 from mature virus particles had no effect on hemifusion but inhibited the next step of pore formation and core entry into the cytoplasm. In contrast several other EFC proteins including A16 and G9 are required for hemifusion. Foo *et al* have previously demonstrated the importance of L1 *N-*myristoylation for complementation of VACV infectivity but did not analyze the entry step [22]. Given that IMP-1088 treatment significantly inhibited L1 *N-*myristoylation by LC-MS/MS proteomics (Figure 2B) western blotting of YnMyr-labeled proteins with an L1 specific antibody was performed to confirm inhibition of L1 *N*-myristoylation. While L1 was pulled down after labeling with YnMyr in untreated cells, there was no detectable L1 pulldown in the presence of IMP-1088 (Figure 6C), providing a direct link between loss of L1 modification and reduction in virus entry.

**Figure 6.**
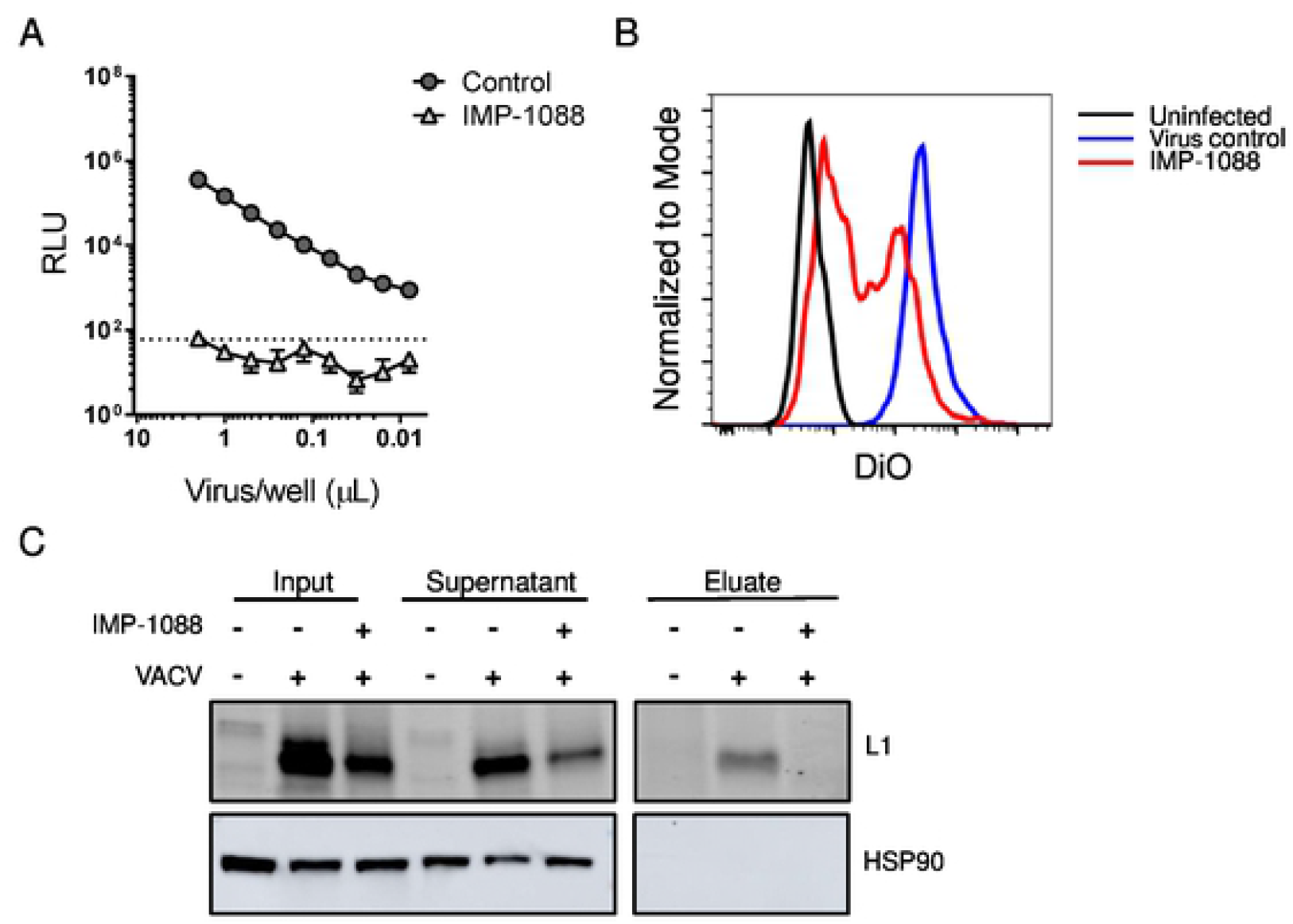
Decrease in VACV infectivity linked to defect in viral entry. (A) IMP-1088 treated virus exhibits lower early gene expression compared to control virus. BSC40 cells were infected with VACV propagated in the absence or presence of 2 μM IMP-1088, and luciferase levels were determined 2 hpi as a surrogate of early protein synthesis. Dotted line indicates level of detection. (B) Lower membrane fusion between cells and IMP-1088 treated virus compared to control virus. Virus grown in the presence or absence of IMP-1088 was purified and labelled with fluorescent dye DiO (same virus as Figure 4). Cells were infected with DiO-labeled virus at RT (to determine background signal) and at 37°C. Transfer of fluorescent dye from viral membrane to cellular membrane was measured by flow cytometry. (C) Western blot to confirm pulldown of L1 protein after metabolic labelling with YnMyr, chemical modification and precipitation using streptavidin resin. The input, supernatant and eluted fractions from uninfected and infected cells in the presence and absence of IMP-1088 were tested for presence of L1 using anti-L1 polyclonal antibody (R180).

### Viruses with G2A mutations of A16 and G9 retain infectivity

The VACV G9 and A16 proteins are essential components of the EFC that have homologs encoded by all poxviruses [25, 26]. Notably, the N-terminal MG *N-*myristoylation motif is conserved in all poxviral G9 and A16 proteins, implying an important role in poxvirus biology. Previous and present biochemical studies show that VACV A16 and G9 are *N-*myristoylated, reinforcing the idea that this modification occurs throughout the poxvirus family. As IMP-1088 reduced *N-*myristoylation of G9 and A16, this effect could contribute to the inhibition of virus entry and infectivity. To investigate this possibility, mutant viruses VACV WR-A16(G2A) and VACV WR-G9(G2A) were constructed. Contrary to our expectation, the mutants remained fully infectious as shown by their relatively unchanged genome/PFU ratios compared to the parental virus (Figure 7A). The mean size of plaques formed by VACV WR-A16(G2A) were considerably smaller and those of VACV WR-G9(G2A) slightly smaller compared to the parental VACV WR, suggesting diminished virus spread (Figure 7B). Further studies are needed to determine the nature of this effect.

**Figure 7.**
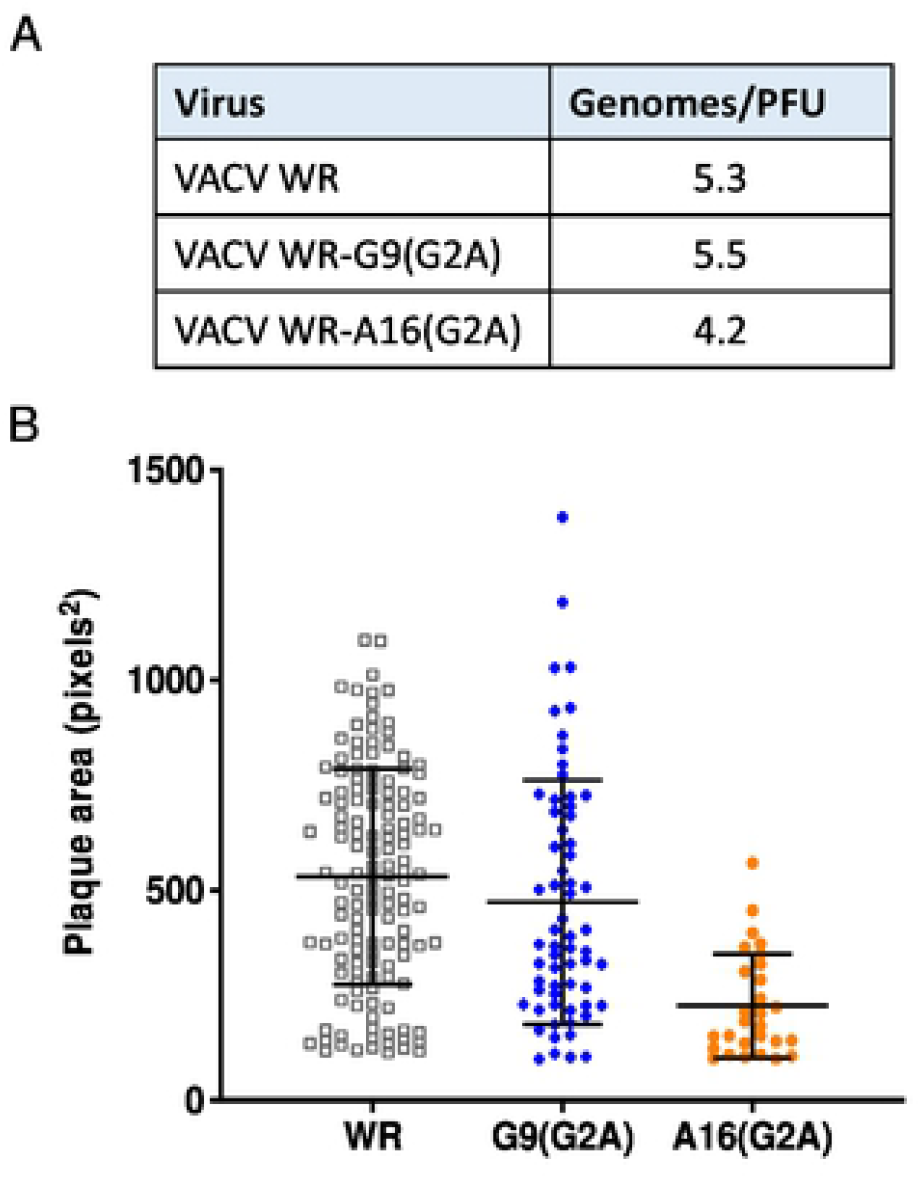
Genomes and infectivity of G9 and A16 viruses with G2A mutations. (A) Ratios of genomes to infectious units. WR, VACV WR-G9(2GA), and VACV WR-A16(G2A) mature virions were purified from infected cells and the infectivity determined by plaque assay. DNA was extracted from the purified virions and genome copies were quantified by ddPCR. (B) Plaque sizes. The areas of plaques formed by purified VACV WR, VACV WR-G9(G2A) and VACV WR-A16(G2A) from a representative experiment are shown. The areas of WR-A16(G2A) plaques were smaller than VACV WR in each of three independent experiments and those of WR-G9(G2A) were similar to those of VACV WR in one experiment and slightly smaller in two others.

## DISCUSSION

Lipidation of VACV-encoded proteins, either with myristic acid or palmitic acid, can be essential for virus infection. A Glycine to Alanine mutation in the N-terminal MG *N*-myristoylation motif of L1 renders VACV deficient in entry [22, 27]. Previous studies putatively identified 6 *N-*myristoylated VACV proteins using radiolabeled analogs [28]. Analysis of VACV encoded proteins by bioinformatics identified 11 proteins with a potential *N-*myristoylation motif (“MG”, at the N-terminus), including the previously confirmed proteins. Five of these were predicted to contain a *N-*myristoylation motif using the program Myristoylator based on additional consensus sequences at the N-terminus [29]. In this study, we used a combined approach of quantitative proteomics and molecular biology to evaluate the impact of a human NMT inhibitor, IMP-1088, on host and VACV protein *N-*myristoylation and viral infection.

IMP-1088 inhibited VACV infection with an EC_50_ concentration of 0.1 μM without detectable cytotoxicity up to 10 μM, at which concentration IMP-1088 is known to entirely inhibit NMT activity in human cells [18]. To determine the NMT protein substrates most strongly impacted by IMP-1088, cells were metabolically labeled with myristic acid analog YnMyr followed by bio-orthogonal ligation, enrichment and quantitative proteomic analysis. Three proteins, namely L1, G9 and A16, were detected. *N*-myristoylation of VACV encoded protein L1 was the most strongly inhibited among all substrates identified in the presence of IMP-1088. Using ^3^H-myristic acid, 4 proteins, L1, A16, G9 and E7 were previously identified as *N-*myristoylated proteins bearing an optimal sequence motif, MGxxxS/T/A/C/N based on bioinformatic analysis [30]. Whilst 11 VACV proteins bear an N-terminal MG motif by sequence analysis, we reliably detected only L1, A16 and G9 by proteomics. Previous mass spectrometric studies of purified viral particles identified the *N-*myristoylation mark in L1 and E7 but not in A16 and G9 likely because of their low abundance [31]. In addition, significant differences in co-translational and post-translational *N*-myristoylation of host proteins was observed, with a reduction in *N*-myristoylation of host proteins detected after VACV infection compared to uninfected controls. An important caveat is that YnMyr was added after VACV infection, and therefore the changes observed in *N*-myristoylated VACV proteins are limited to newly translated proteins. The generalized shut down of host protein synthesis, which occurs following VACV infection [23], rather than a decrease in *N*-myristoyl transferase activity may account for the observed reduction in labeled host proteins.

Immediately following VACV entry and the release of viral cores into the cytoplasm, early mRNAs are transcribed and translated. After early gene expression, uncoating of viral cores releases encapsidated DNA which leads to DNA replication and the expression of intermediate and late genes [32]. IMP-1088 potently decreased VACV infection and spread, however it did not significantly affect viral early or late protein synthesis suggesting no effect on DNA replication and gene expression. By electron microscopy, the progression of VACV morphogenesis from crescent to extracellular virus was observed in the presence of IMP-1088 indicating that virus formation was not abrogated. However, purified viral particles generated in the presence of IMP-1088 were found to be less infectious than virus from untreated infected cells, and exhibited reduced membrane fusion and core release in the cytoplasm. This overall phenotype is characteristic of entry-deficient viruses that lack functional entry/fusion complex (EFC) proteins, including L1 [33]. Our mass spectrometry analyses revealed L1 as the VACV protein with the most significant change in *N*-myristoylation in the presence of IMP-1088. Additionally, the absence of L1 in YnMyr pull down fractions comparing IMP-1088 treated and untreated cells demonstrated a defect in YnMyr incorporation by L1 in the presence of IMP-1088. Nevertheless, YnMyr incorporation into two additional *N-*myristoylated EFC proteins, A16 and G9, was also inhibited by IMP-1088. To investigate the role of *N-*myristoylation of these proteins, we constructed mutants with G2A mutations in the essential MG motif. The A16 G2A and G9-G2A mutants retained infectivity although the size of plaques formed by VACV with the A16 G2A mutation were smaller than those of the parental virus. Based on the retained infectivity of A16 and G9 mutant viruses in our study, and the previous report showing the lack of complementation of L1 G2A mutant gene [22], L1 appears to play the most critical role in the IMP-1088 mediated inhibition of VACV infection. Exactly how *N-*myristoylation of L1 enables virus entry is not known, though it has been shown that this modification is necessary for formation of intramolecular disulfide bonds in L1 [22, 33].

Due to the importance of L1 for virus entry, it is targeted for antibody mediated neutralization and protein subunit-based vaccines against VACV [34-36]. Recently, two independent monoclonal antibody cocktails containing an anti-L1 VACV neutralizing antibody conferred protection *in vivo* against VACV and monkeypox virus when given prophylactically, further emphasizing its indispensable role in poxviral pathogenesis [37, 38]. However, anti-L1 mAb cannot neutralize EV particles, and protection against OPXV infection *in vivo* requires combination with mAbs that target EV proteins[39]. Even though the EV membrane dissociates prior to entry to expose the MV membrane, epitopes recognized by anti-L1 antibodies may remain inaccessible due to protection from ruptured membranes or spatial separation of fusion activity [40-43]. In our study, we demonstrate that the NMT inhibitor IMP-1088 potently inhibits both MV and EV particles by preventing *N*-myristoylation of L1. Given the high genetic identity between VACV and VARV L1 homologs (99.2%), additional studies that evaluate the *in vivo* efficacy of IMP-1088 against VACV as well as its *in vitro* potential against VARV are warranted. As IMP-1088 targets a different step in the OPXV life cycle than tecovirimat (TPOXX®, EV formation) and brincidofovir (TEMEXA®, DNA replication), the two FDA approved therapeutics [44, 45], the possibility of synergistic effects of combination therapy with both compounds may also warrant further investigation.

## MATERIALS AND METHODS

### Viruses and cell lines

The viruses used in this study have been previously described and include VACV WR-GFP [46], VACV WR-A4FP [47], VACV WR-pE/L-LUC [48], VACV WR-pF17R-LUC [49] and VACV IHDJ. VACV WR-G9(G2A) and VACV WR-A16(G2A) were constructed by homologous recombination and contain an adjacent GFP or dsRed gene, respectively, regulated by the VACV P11 promoter, which was used for selection as previously described [25, 26]. The codon changes in the mutated genes were confirmed by Sanger sequencing. Virus stocks were grown in BS-C-40 or BS-C-1 cells with DMEM containing 2% (v/v) fetal bovine serum (FBS) and stored at -80 °C prior to use. BS-C-40, BS-C-1 and HeLa cell lines were passaged in DMEM with 10% (v/v) FBS and 10 units/mL of penicillin and 100 μg/mL of streptomycin. For all experiments involving viral infection, DMEM containing 2% (v/v) FBS was used as a diluent or culture media.

### Virus yield quantification

To quantify total virus yield, HeLa cells were infected with VACV in the presence or absence of IMP-1088 for 24 h at 37 °C. The following day, cells and supernatants were collected and frozen at -80 °C. The samples were serially diluted and titrated by plaque assay using BSC40 cells (described below). To specifically quantify EV yield, only the supernatants of infected cells were collected, and subsequently titrated by plaque assay (described below).

### Plaque assay

BSC40 cells were infected with serially diluted VACV-containing samples for 1 h at room temperature (RT). The infected cells were washed three times to remove unbound virus. Each dilution was tested in triplicate. An overlay containing 2% (w/v) methylcellulose was added to each well, and plates were incubated at 37 °C for 72 h. Monolayers were then fixed and stained using crystal violet stain containing 10% (v/v) formalin. Plaques were counted and used to determine viral titer in the presence or absence of treatments. Areas of individual plaques formed in BS-C-1 cells and stained with crystal violet were determined using an EVOS cell imaging system (Thermo Fisher Scientific).

### Viral spread assay

The viral spread assay was performed as previously described [50]. Briefly, IMP-1088 was serially diluted and mixed with VACV-WR-GFP virus diluted in DMEM-2. The mixture was used to infect HeLa cells seeded in 96-well plates (Corning, 06-443-2) at 37 °C for 24 h. A cytosine arabinoside (AraC; 40 μg/mL) treatment and other controls were included on each plate, and the IMP-1088 dilution series was tested in duplicate. After 24 h, the cells were fixed with 4% (w/v) paraformaldehyde for 15 min at RT and stained with 4’,6-diamidino-2-phenylindole (DAPI) for nuclei visualization for 10 min at RT. The plates were imaged using the ArrayScan XTI High Content Screen (HCS) reader, and the percentage of GFP and DAPI positive cells was quantified using the HCS Studio Cell Analysis software as previously described [51]. Raw data was analyzed using GraphPad Prism software (GraphPad Software, v7) to determine the concentrations required for 50% inhibition (EC_50_) of viral spread relative to the no treatment control.

### LDH Cytotoxicity Assay

The LDH Cytotoxicity Assay was performed using the LDH Cytotoxicity Assay kit (Thermo Scientific Pierce, 88953) as per the manufacturer’s instructions. Serially diluted compound (10-0.62 μM) was added to HeLa cells and incubated at 37 °C for 24 hrs. Supernatants were collected and the levels of extracellular lactate dehydrogenase (LDH) expressed in the supernatants were quantified. Data was analyzed using GraphPad Prism software (GraphPad Software, v7).

### Early/Late protein and Late protein synthesis

BSC40 cells were infected with either a VACV WR-pE/L-LUC which expresses luciferase by a synthetic early/late gene promoter or with VACV WR-pF17R-LUC, which expresses luciferase under a F17R gene late promoter. Cells were infected with virus at MOI 3 for 1 h at RT. After infection, cells were washed three times with 1X PBS to remove unbound virus. IMP-1088 diluted to 10 μM was added to cells and incubated at 37 °C for either 2 h or 24 h to quantitate early/late or late protein synthesis respectively. Cells were lysed with Reporter Lysis Buffer, subjected to a freeze-thaw cycle to lyse cells and luciferase activity was measured using the Luciferase Assay System (Promega, Madison, WI) according to manufacturer’s instructions. Luciferase activity was measured using an ENSPIRE plate reader (PerkinElmer, Waltham, MA, United States).

### Electron Microscopy

BS-C-40 cells were infected with VACV IHDJ in the presence or absence of 2 μM IMP-1088 for 24 h at 37 °C. Following infection, cell monolayers were gently scraped, pelleted, and processed for thin-section electron microscopy (EM). Specimens were fixed in buffered 2.5% (w/v) glutaraldehyde, fixed in 1% (w/v) osmium tetroxide, stained *en bloc* with 4% (w/v) uranyl acetate, dehydrated through a graded alcohol and acetone series, and embedded in a mixture of Epon substitute and Araldite epoxy resins. Thin sections were stained with 4% (w/v) uranyl acetate and Reynolds’ lead citrate [52].

### Purification and specific infectivity of VACV

BSC-40 cells were infected with VACV WR-pE/L-LUC virus in the presence or absence of 2 μM IMP-1088 for 24 h at 37 °C. The cells were harvested, and viruses were purified using a sucrose cushion followed by sucrose gradient centrifugation and resuspended in equal volumes. Purified virus stocks were frozen at -80 °C until use. To assess the relative number of viral particles present in the untreated and IMP-1088-treated stocks, we measured viral protein content and the quantity of genomic viral DNA using SD*S-*PAGE followed by Coomassie staining and Real-time PCR, respectively. Thereafter, the viruses were titrated by plaque assay to determine infectious virus yield or plaque forming unit (pfu) of untreated and IMP-1088-treated stocks.

### Quantitative PCR

Viral DNA samples were tested with a quantitative orthopoxvirus generic PCR assay based on TaqMan® chemistry and technology described elsewhere [53]. Each 20 μL reaction contained 6.5 μL of RNase/DNase free water (Clonetech, Mountainview, CA), 10 μL of TaqMan® Fast Advanced Master Mix (Life Technologies, Grand Island, NY), 0.5 μL each of forward and reverse primer at 50 μM and 0.5 μL of probe at 10 μM, to which 5 μL of DNA were added. Cycling parameters for the real-time PCR were 95 °C for 20 s, followed by 40 cycles of 95 °C for 3 s and 60 °C for 30 s performed on a 7500 Fast Dx Real-Time PCR Instrument (Life Technologies, Grand Island, NY).

### Genome/PFU ratios

VACV mature virions were purified from infected cell homogenates by sedimentation through a sucrose cushion, and Infectivity was determined by plaque assay. The purified virions were treated with Benzonase (Sigma, ST. Louis) to remove adventitious DNA after which genomic DNA was extracted and quantified by digital droplet PCR as described [54].

### Virus entry assay

BSC40 cells were infected, for 2 h at 37 °C, with serially diluted VACV WR-pE/L-LUC virus grown in the presence of IMP-1088 or DMSO. The cells were subsequently lysed using the Reporter Lysis Buffer and freeze-thawed once, and luciferase activity was measured using the Luciferase Assay System according to manufacturer’s instructions. Luciferase activity was measured using an ENSPIRE plate reader.

### Viral fusion assay

To evaluate membrane fusion of VACV virus, both viruses were labeled with lipophilic tracer DiO as previously described [55]. Briefly, purified untreated VACV-WR and IMP-1088-treated VACV-WR were incubated with DiO diluted in 1X PBS (Thermofisher) in the dark for 20 min at RT. The viruses were subsequently washed and pelleted three times to remove unbound DiO. BSC40 cells were incubated with each of the DiO-labeled viruses for 1 h at RT to bind, and then washed three times with 1X PBS to remove unbound virus. The virus + cell mixture was then incubated at 37 °C for 90 min to allow membrane fusion. Following fusion, the samples were fixed with 4% (w/v) paraformaldehyde for 15 min and then analyzed by flow cytometry using the Attune Nxt instrument to determine percent DiO^+^ cells.

### Metabolic tagging and chemical proteomics to identify *N-*myristoylated proteins

In triplicate, HeLa cells were grown in T75 flasks to 80-90% confluency, followed by infection with VACV-Luc as well as a metabolic tagging pulse of 20 μM tetradec-13-ynoic acid (YnMyr) for 24 h, in the presence or absence of 2 μ? IMP-1088. In parallel, HeLa cells were treated identically but in absence of VACV infection, as uninfected controls. Then, the cells were washed with PBS and lysed by scraping in lysis buffer (1% (v/v) Triton X-100, 0.1% (w/v) SDS and EDTA-free protease inhibitor cocktail (Roche, 11873580001) in PBS, pH 7.4). Proteins were precipitated (methanol:chloroform:water at 4:1:2), the pellet washed with cold methanol and resuspended in 1% (v/v) Triton X-100, 0.1% (w/v) SDS and EDTA-free protease inhibitor cocktail (Roche, 11873580001) in PBS, pH 7.4. Protein concentrations were determined (BCA assay kit, Thermo Fisher 23250). Lysates (300 μg total protein) were incubated with premixed copper-catalyzed cycloaddition (CuAAC) ligation reagents (100 μM AzRB, 1 mM CuSO_4_, 1 mM TCEP (Sigma C4706), and 100 μM TBTA (Sigma 678937)) while vortexing for 1 h at RT. After quenching with 10 mM EDTA, proteins were precipitated (methanol:chloroform:water at 4:1:2), the pellet washed with cold methanol and resuspended in 0.2% (w/v) SDS in 50 mM HEPES (Sigma 54457), pH 8.0. After enrichment on NeutrAvidin-coupled agarose beads and stringent detergent washing, enriched proteins were digested on-bead with trypsin (Promega, V5111). Peptides were acidified with 0.5% (v/v) TFA (Sigma T6508), desalted on Stage Tips, and analysed by Label Free Quantification by nanoLC-MS/MS on a Thermo QExactive instrument as described previously [12, 19]. The proteomic analysis data was processed using MaxQuant v1.6.4.0 with built-in Andromeda search engine [56]. Peptides were identified from the MS/MS spectra by searching against the human (UP000005640) and the VACV virus (UP000000344) FASTA proteome references with both canonical and isoforms. Cysteine carbamidomethylation was set as a fixed modification and methionine oxidation, N-terminal acetylation and YnMyr-AzRB2 as variable modifications. ‘Trypsin/P’ was the digestion mode enzyme, and up to two missed cleavages were allowed. The ‘match between run’ option was selected, along with ‘unique and razor peptides’ for protein quantification. Processed data were further analyzed using Perseus v1.6.2.3, RStudio 1.4.1106 (R version 4.0.4) and GraphPad Prism v8.0. Prerequisite for statistical significance testing was a minimum of two identifications per triplicate. To determine statistical significance within Volcano plots, 250 randomized, two-sided Student t-tests were performed with false discovery rate (FDR) set to 0.05 and S0 to 0.1, and a minimum of 2 unique peptides per protein were required.

Lysates (300 μg total protein) were incubated with premixed copper-catalyzed cycloaddition (CuAAC) ligation reagents (100 μM AzRB, 1 mM CuSO_4_, 1 mM TCEP (Sigma C4706), and 100 μM TBTA (Sigma 678937)) while vortexing for 1 h at RT. After quenching with 10 mM EDTA, proteins were precipitated (methanol:chloroform:water at 4:1:2), the pellet washed with cold methanol and resuspended in 0.2% (w/v) SDS in 50 mM HEPES (Sigma 54457), pH 8.0. After enrichment on NeutrAvidin-coupled agarose beads and stringent detergent washing,

### Validation of *N*-myristoylation of VACV L1 protein

Samples were prepared as described in previous section [11]. Briefly, 50 μg proteins were incubated with premixed CuAAC ligation reagents [100 μM AzRB, 1 mM CuSO_4_, 1 mM TCEP (Sigma C4706), and 100 μM TBTA (Sigma 678937)] and vortexed for 1 h at RT. After quenching with 5 mM EDTA, proteins were precipitated (methanol:chloroform:water at 4:1:2), the pellet washed with cold methanol and resuspended in in PBS containing 2% (w/v) SDS and 10 mM DTT. Samples were centrifuged at 17,000 g for 10 min and an aliquot was set aside as the “input” sample. The remaining volume was submitted for biotin-enrichment on magnetic Streptavidin beads (NEB S1420S), by vortexing for 90 min at RT. The mixture was separated by placing the tube on a Magna GrIP rack magnet (Sigma, 20-400) for 2 min, allowing collection of the supernatant, and stringent detergent washing of the beads. The beads were finally boiled in Laemmli sample buffer (10% (w/v) SDS, 25% (v/v) β-mercaptoethanol and 0.02% (w/v) bromophenol blue in 50% (v/v) glycerol in 1 M Tris*-*HCl, pH 6.8) for 10 min at 95 °C. Input and supernatant samples were boiled at 95 °C for 5 min. To detect L1 protein, samples were loaded and run on a 12% (w/v) SD*S-*PAGE gel, transferred onto a nitrocellulose membrane by blotting and incubated for 1 h with 5% (w/v) non-fat milk in PBS supplemented with 0.1% (v/v) Tween20. The membrane was incubated with an anti-L1 antibody (R180) [22] overnight and with a secondary rabbit antibody IRDye 800CW (Li-Cor 926-32211) for 1 h. The blots were imaged using the Odyssey CLx (LI-COR Biosciences).

## ACKNOWLEDGEMENTS

The study was supported by the CDC Intramural Research and Biomedical Advanced Research and Developmental Authority (BARDA) and an ORISE Postdoctoral Fellowship (LP). The European Commission (Marie Sklodowska Curie Individual Fellowship grant 752165 to WWK), EPSRC (Impact Acceleration Account grant PS1042 to WWK and EWT), Cancer Research UK (C29637/A21451 and C29637/A20183 to E.W.T.). Work in the E.W.T. laboratories is supported by the Francis Crick Institute, which receives its core funding from Cancer Research UK (FC001057 and FC001097), the UK Medical Research Council (FC001057 and FC001097), and the Wellcome Trust (FC001057 and FC001097). Partial support was provided by the Division of Intramural Research, NIAID. The findings and conclusions in this report are those of the authors and do not necessarily represent the official position of the Centers for Disease Control and Prevention.

## AUTHOR CONTRIBUTIONS

LP and WWK contributed to the design of the viral and proteomic studies, performed experiments, analyzed data and co-wrote the manuscript.

MF, KW, CSG, CAC, SO performed experiments and analyzed data.

RS and BM contributed to the design of the studies and review of the data.

EWT contributed to direction and design of the proteomic studies, co-wrote the manuscript and co-led the study.

SP contributed to direction and design of the viral studies, co-wrote the manuscript and co-led the study.

## Supplementary data

**Supplementary Table 1.**
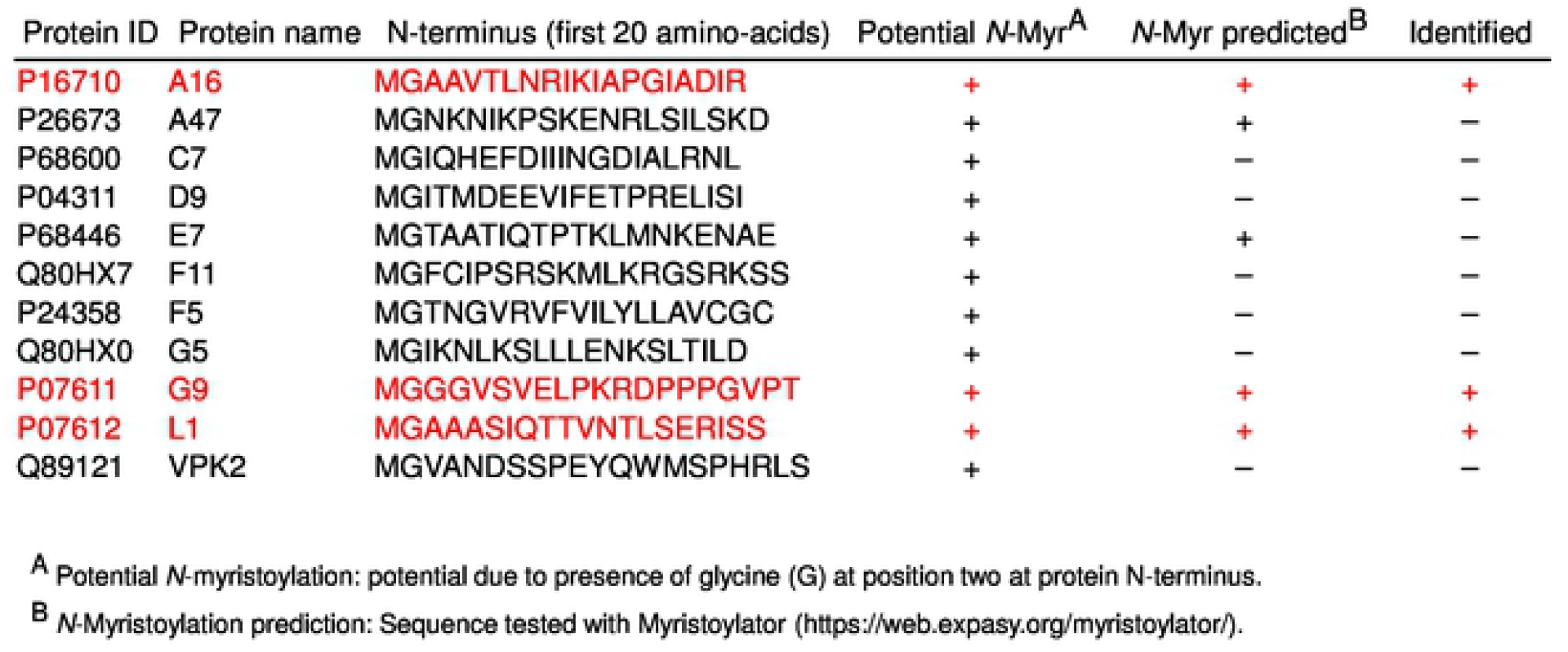
*N*-myristoylated proteins present in VACV proteome. Validated *N*-myristoylated proteins A16, G2 and L1 shown in red, and potentially *N*-myristoylated proteins shown in black.

**Supplementary Figure 1.**
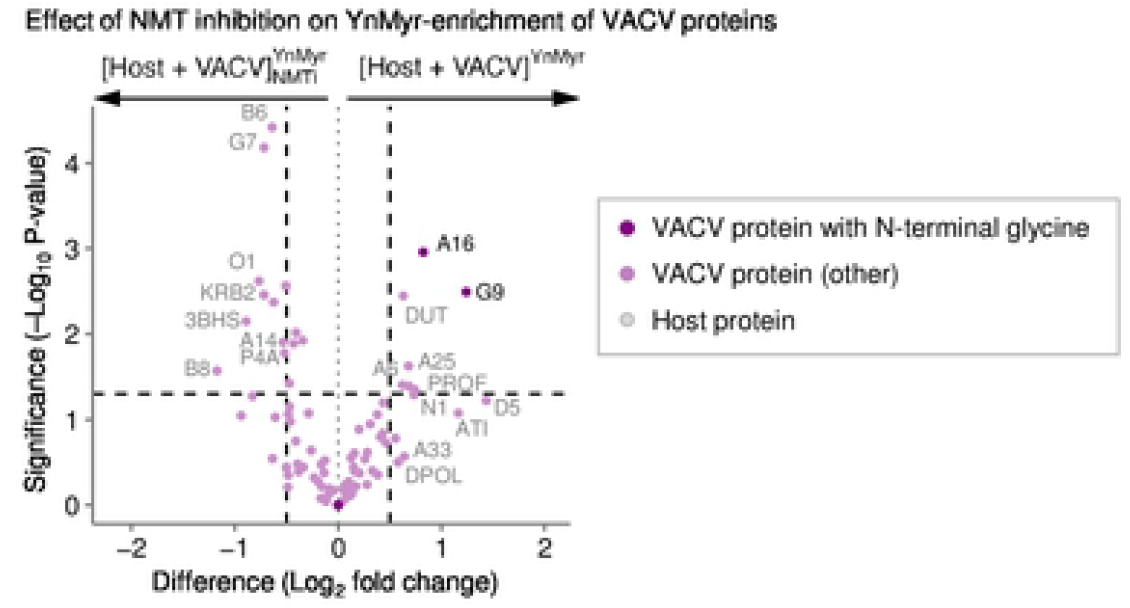
Chemical proteomic identification of *N*-myristoylated VACV proteins. Target engagement of NMT inhibitor (NMTi = IMP-1088) on VACV proteins in infected host cells. Left vertical line depicts -0.5 Log_2_ fold change, right of +0.5; horizontal line depicts significance cut-off (P = 0.05); 3 *N*-myristoylated VACV proteins depicted in purple, 115 VACV proteins in pink, 2738 human proteins in gray.

**Supplementary Figure 2.**
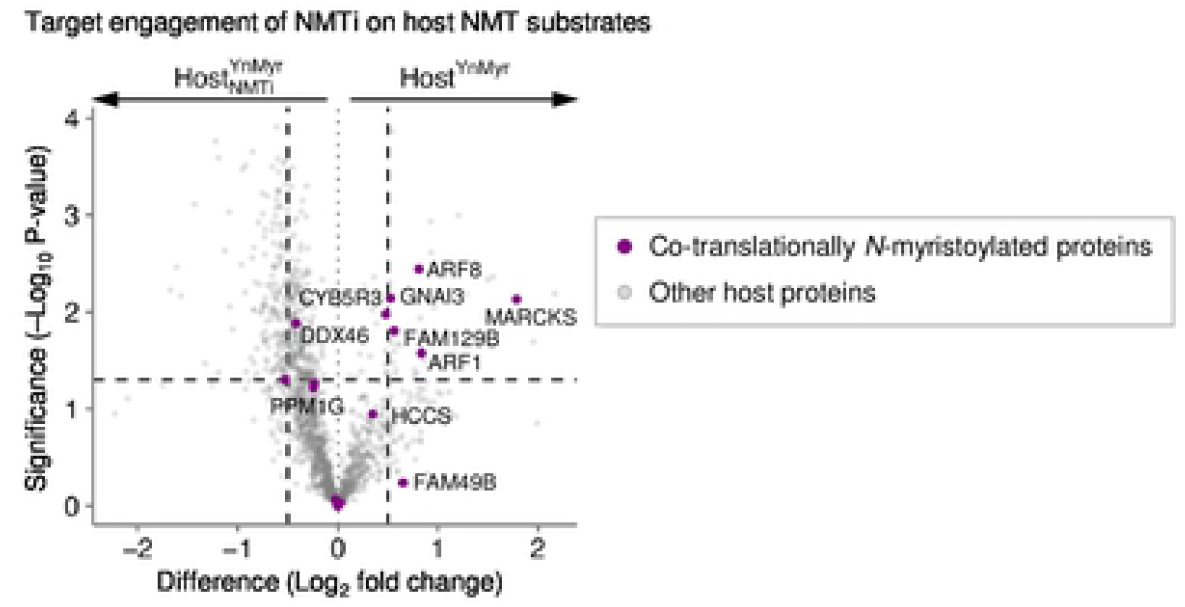
Target engagement of NMTi on co-translationally *N*-myristoylated host proteins. Target engagement of NMT inhibitor (NMTi = IMP-1088) in host cells, as visualized by the effect on 32 known co-translationally *N-*myristoylated proteins of the host (purple). Left vertical line depicts - 0.5 Log_2_ fold change, right of +0.5; horizontal line depicts significance cut-off (P = 0.05). Other proteins in gray.

**Supplementary Figure 3.**
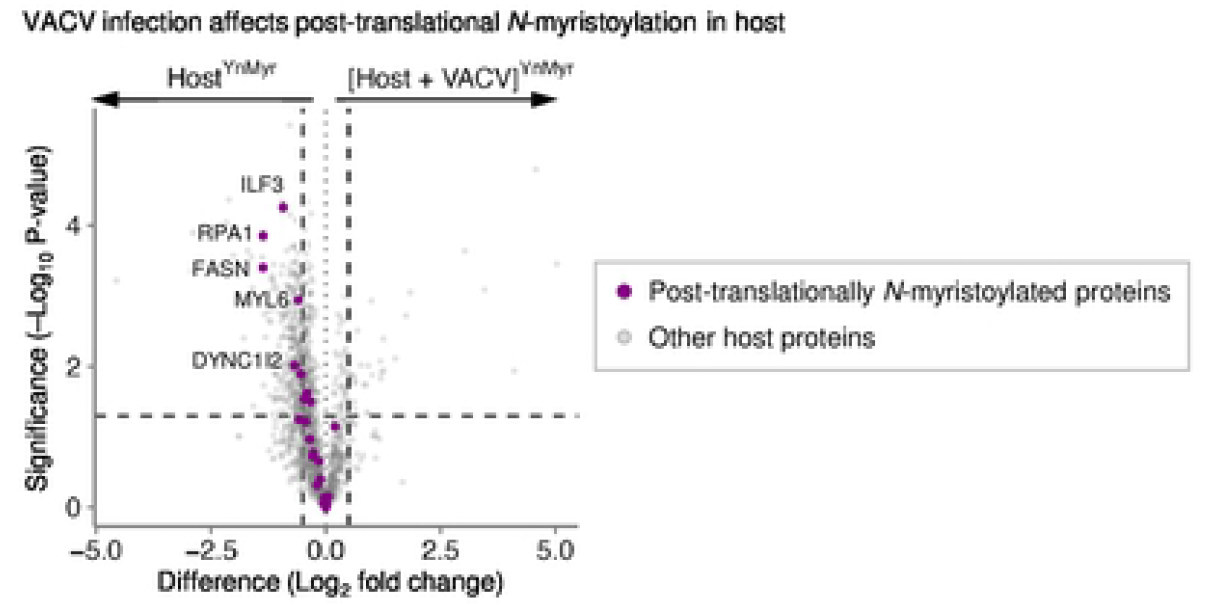
VACV reduces abundance post-translationally *N*-myristoylated proteins of host. Effect of VACV infection on 27 known post-translationally *N-*myristoylated protein levels of the host (purple). Left vertical line depicts -0.5 Log_2_ fold change, right of +0.5; horizontal line depicts significance cut-off (P = 0.05). Other proteins in gray.

